# Bumblebees navigate using path integration while walking

**DOI:** 10.1101/2022.03.02.482643

**Authors:** Rickesh N. Patel, Julian Kempenaers, Stanley Heinze

## Abstract

Path integration is a computational strategy that allows an animal to maintain an internal estimate of its position relative to a point of origin. Many species use path integration to navigate back to specific locations, typically their homes, after lengthy and convoluted excursions. Hymenopteran insects are impressive path integrators, directly returning to their hives after hundreds of meters of outward travel. Recent neurobiological insights have established hypotheses for how path integration may be mediated by the brains of bees, but clear ways to test these hypotheses in the laboratory are currently unavailable. Here we report that the bumblebee, *Bombus terrestris*, uses path integration while walking over short distances in an indoor arena. They estimate accurate vector distances after displacement and orient by artificial celestial cues. Walking bumblebees also exhibited systematic search patterns when home vectors failed to lead them accurately back to the nest, closely resembling searches performed by other species in natural conditions. We thus provide a robust experimental system to test navigation behavior in the laboratory that reflects most aspects of natural path integration. Importantly, we established this assay in an animal that is both readily available and resilient to invasive manipulations. In the future, our behavioral assay therefore can be combined with current electrophysiological techniques, opening a path towards directly probing the neural basis of the sophisticated vector navigation abilities of bees.

## INTRODUCTION

Path integration is a computational strategy that allows animals to maintain an internal estimate of their position within the environment with respect to a point of origin. Many species use path integration for navigation, particularly when they return to a location of interest after travelling away from it. During path integration, an animal monitors its angular and translational displacements to continually compute the distance and angle to the origin of its excursion, representing the most direct path back to the reference point, the home vector. This home vector can guide steering movements to enable the animal to return to its original location^1^.

The use of path integration has been most clearly demonstrated in central place foraging animals that return to a home location between foraging bouts; examples include insects^2–4^, crustaceans^5,6^, and mammals^7,8^. Additionally, a spider^9^ and a fruitfly^10^ have been shown to use path integration to return to a food source. However, path integration has been most thoroughly studied in hymenopterans, including desert ants^11^ and honeybees^12^, champion path integrators who are able to accurately locate their nests/hives after hundreds of meters of tortuous travel.

To update their positional estimate, hymenopteran insects primarily rely on external cues (allothetic cues) as sources for directional information and self-generated movement cues (idiothetic cues) for distance information. While many cues can be used to obtain directional information, celestial cues are among the most prevalent. Many species, including hymenopterans, primarily orient using the skylight polarization pattern^6,12–17^, created by the scattering of sunlight in the Earth’s atmosphere, and the solar azimuth^6,18–22^. For measuring distance, honeybees use optic flow^23,24^, the pattern of apparent motion of the visual scene when moving, whereas desert ants primarily use a stride integrator^25,26^.

Although path integration is a robust strategy, positional estimates obtained using idiothetic information accumulate error and therefore, are not perfect. Responding to this error, path-integrating animals characteristically initiate a stereotyped search behavior if they fail to reach their goal after travelling the distance indicated by their home vectors. These patterns are generally composed of loops that are centered around the initiation point of the search and increase in size over time, a highly effective strategy for locating a goal in the absence of a positional marker^3–5,27–29^.

Like honeybees, bumblebees (*Bombus terrestris*) forage at flower patches for pollen and nectar before returning to their nests. This species has become an important model for hymenopteran research^30–32^ due to its relativity small, manageable hive size, commercial availability, relatively large body size, and robustness to invasive experimental procedures. All these characteristics are key assets to experimental investigations in the laboratory, including those aimed at unravelling the still enigmatic neural basis of path integration. However, while bumblebees are closely related to honeybees and therefore likely use path integration during navigation as well, this ability has to date not been unambiguously demonstrated in this species.

Recent gains in understanding the neural circuitry that likely underlies path integration in bees has yielded a region of the arthropod brain termed the central complex as the main cellular substrate for path integration and general navigation control. This progress is based on comparisons between detailed electron-microscopical maps of the central complexes of flies^33^ and bumblebees^34^, combined with physiological data and anatomically constrained computational models in a sweat bee^35^. However, to link neural circuits of the central complex to the sophisticated path integration abilities of hymenopteran insects, circuit data need to be combined with functional and behavioral data in a single model species. Despite the lack of behavioral data but given their robustness for electrophysiology and our advanced understanding of their central complex circuits, we propose bumblebees as the prime model for this endeavor.

Here, we developed a robust behavioral assay demonstrating path integration in bumblebees. By exploiting walking behavior on a flat surface over relatively short distances, using artificial skylight cues as sole sources of directional information, our assay allows examination of hymenopteran vector navigation systematically in a controlled setting. Importantly, the assay is also highly amenable to electrophysiological experiments and other invasive approaches. We thus not only demonstrate robust path integration abilities in walking bumblebees, but also offer the chance to further dissect complex vector navigation at the behavioral as well as neural level-within a single insect species.

## RESULTS

### Bumblebees maintain an active hive while walking in the laboratory

First, we asked whether a bumblebee hive consisting entirely of walking bees would maintain a steady population of foraging individuals that can be probed for navigational abilities. We thus allowed a hive of bumblebees with clipped wings to forage in a featureless circular arena 1.5 m in diameter. The arena and a hive placed under it were joined by a tube, so that the entrance was hidden from view while the bees were foraging. Equally hidden from view was a conical feeder, which was filled with pollen and sugar solution and placed at the center of the arena (Figures 1 and S1). Illumination was provided by an overhead light source consisting of white and ultraviolet LEDs and a composite filter constructed of a polarizer and a diffuser (Figures 1A, C). This light source could either deliver a linearly polarized light field, as was the case during the initial observations of the hive, or a depolarized light field (Figure 1D). Foraging paths to and from the feeder were video recorded from above (Figure 1).

**Figure 1.**
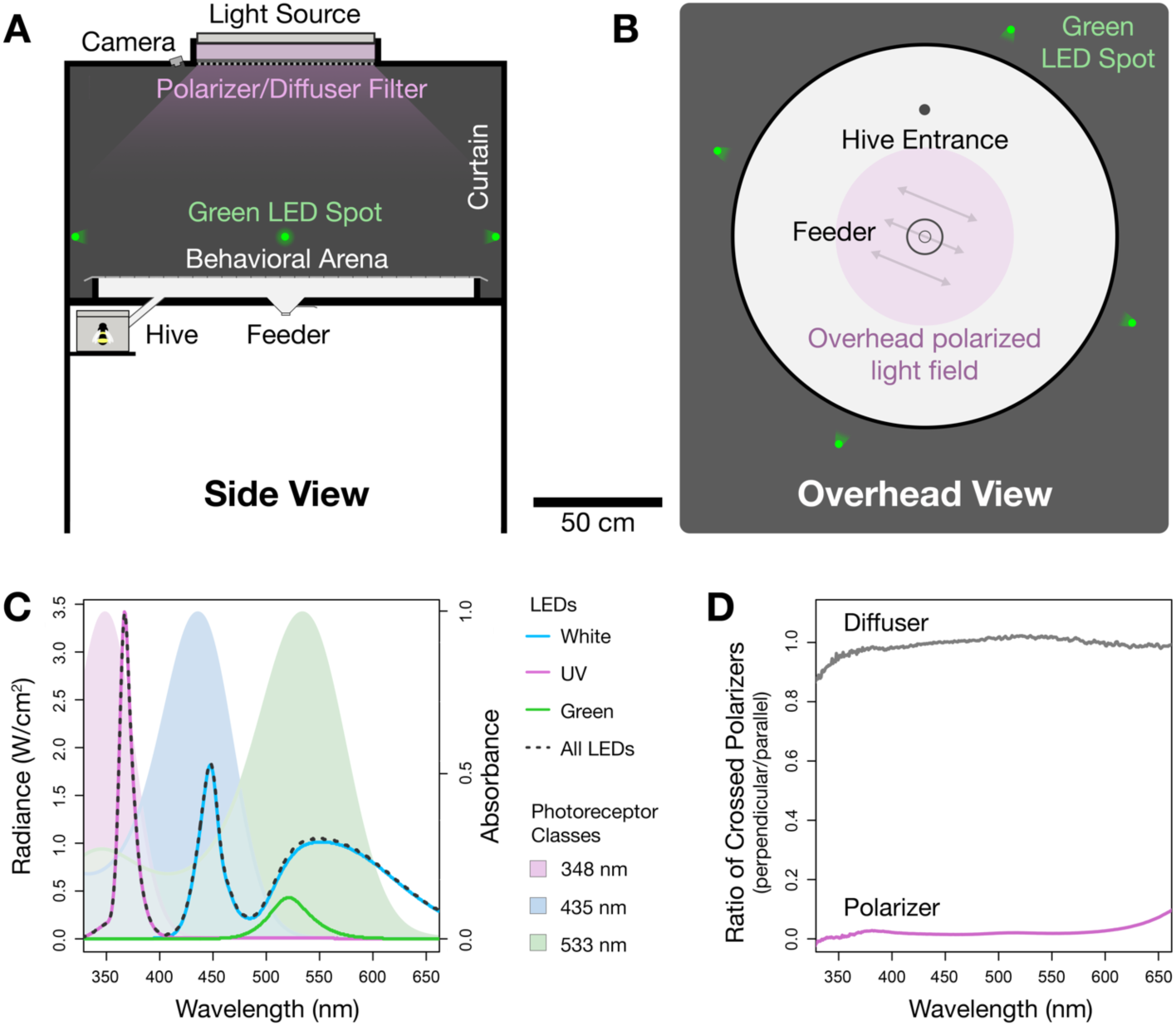
Navigation arena design. **(A)** Side view. **(B)** Overhead view. The circular arena (150 cm diameter) was directly connected to a hive via an entrance located 25 cm from the arena’s periphery (filled circle in B). Sugar-water and pollen was provided in a conical feeder at the arena center. Neither the feeder nor hive-entrance were visible to bees in the arena. The arena was illuminated by an overhead light source that could either be highly polarized or depolarized. Four green point sources were positioned at 90° intervals around the arena. Experiments were video recorded from above. **(C)** Irradiance spectra of light available at the center of the arena (dashed line). Violet and blue curves: irradiance of the isolated UV and white LEDs from the overhead light source. Green curve: irradiance of the green LED point source (at distance of 20 cm). Curves with colored area under the curve: Absorbance spectra of photoreceptor classes in the eyes of *Bombus terrestris dalmatinus* (from Skorupski et al.^37^). **(D)** Ratio of light transmitted through the overhead filter and a second crossed polarizer (perpendicular orientation/parallel orientation) when either the polarizer side (violet) or diffuser side (grey) faced towards the arena. Also see Figure S1.

Over a period of 22 days, initial observations showed that the hive’s foragers were actively making walking foraging trips throughout the entire 12-hour light period of the day (Figure 2A). Peak activity periods of individuals occurred at different times of the day from one another (Figure 2B), and the aging hive actively produce cohorts of new bees who successively took over the foraging duties of older bees (Figure 2C). Thus, despite the unnatural state of the hive and the artificial environment, our hives of walking bees appeared healthy, and the collective activity of the hive largely resembled that reported for hives of flying bees placed in large greenhouses^36^.

**Figure 2.**
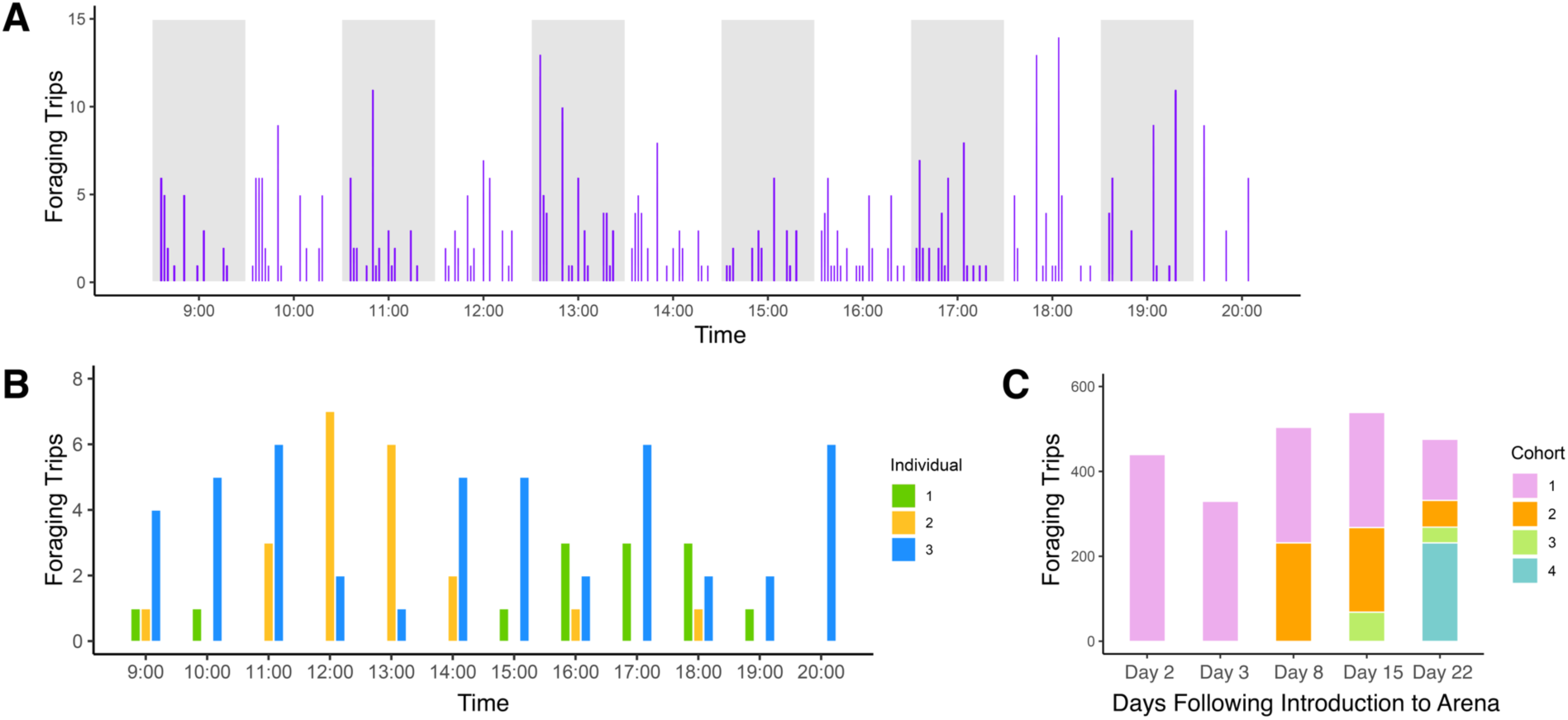
Foraging activity of a hive of walking *Bombus terrestris* in the arena. **(A)** Activity over the course of one representative day light period (day two after introduction to the arena); bees were active throughout the entire light period (9:00-21:00). Alternating grey and white columns: hourly intervals. Violet bars: number of times a single individual made a foraging trip from the hive to the feeder and back again over the course of the corresponding one-hour interval. **(B)** Individuals are active at different times of the day. Colored bars: number of times a single individual made a foraging trip from the hive to the feeder and back again over the course of the corresponding one-hour interval. **(C)** The hive actively produces new individuals that replace older foragers. Colored bars: number of forging trips made by individuals of each cohort per day. Individuals that were new during an observation day were assigned to a new cohort.

### Bumblebees navigate using path integration over short distances while walking

Next, we asked whether walking bumblebees would use path integration to return home after foraging at the feeder. After three days of acclimation to the arena with the overhead polarization field present, bees were released individually from the hive and allowed to locate the feeder. After feeding, bees executed a generally well-directed homeward path, either towards or opposite to the hive’s location (Figure 3B and Video S1). This bimodal orientation is expected since a polarization pattern used as a sole orientation cue indicates only an axis of orientation rather than an absolute direction. If the hive was not encountered at the end of the homeward path, the bees initiated a search behavior, suggesting that *B. terrestris* may use path integration to locate its hive while walking over short distances.

**Figure 3.**
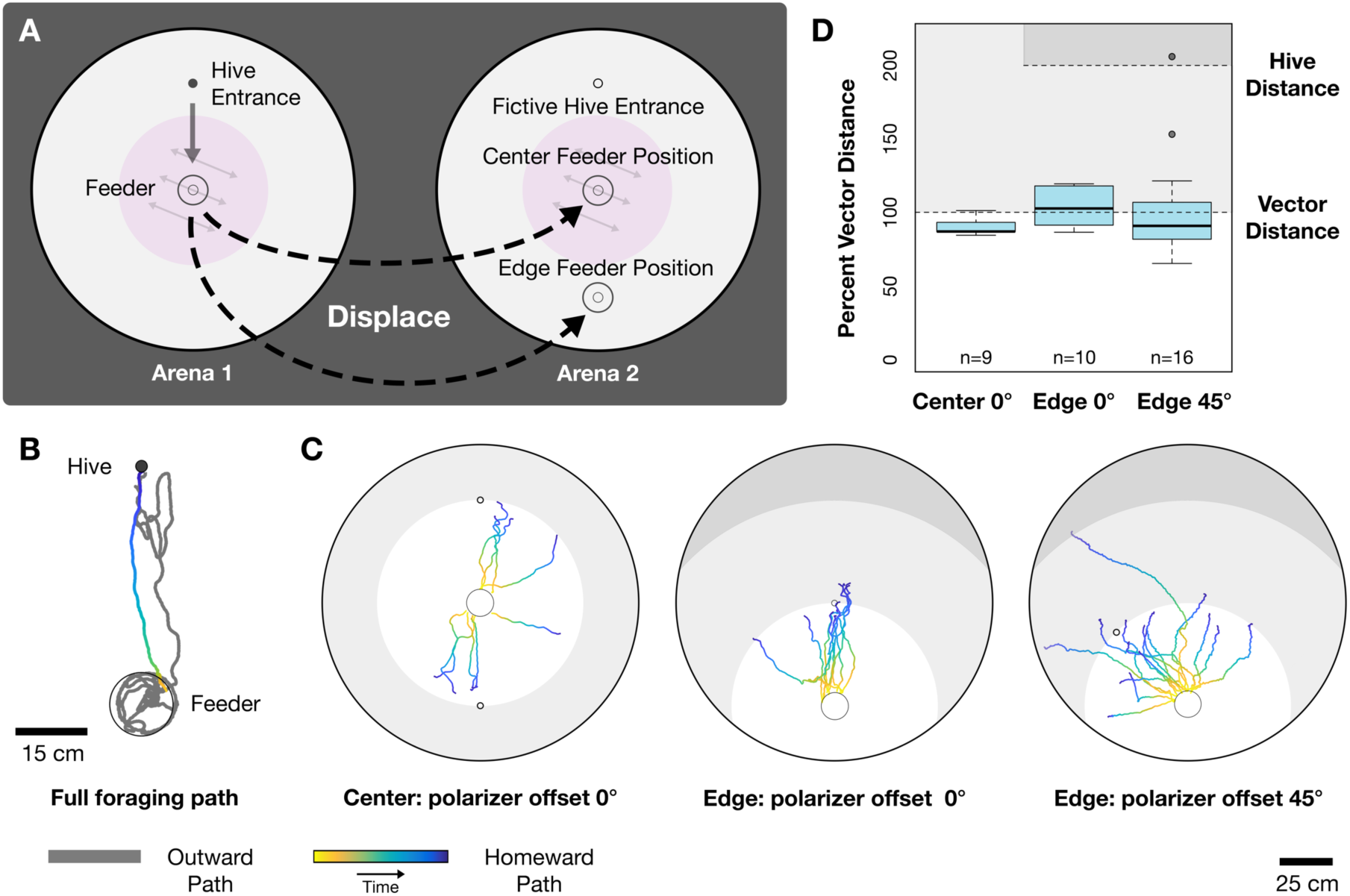
Walking *Bombus terrestris* use path integration to navigate back to the hive while foraging. **(A)** Experimental design: After finding the feeder, bees were trapped and displaced (dashed arrows) to the center or the edge of a second, identical arena that lacked the hive entrance. If the bees use path integration while homing in the arena, they should, after displacement, travel a length equal to the distance between the feeder and the hive entrance in the first arena (vector distance). **(B)** Example of a foraging path from and to the hive when an animal was not displaced. **(C)** Path tracings from all homeward paths before search behaviors were initiated during the three displacement conditions. Shades of grey: regions of the arena further away from the release point than the vector distance (light grey) and further than the hive distance (dark grey, only for edge-displacements). For center-displacements, hive distance and vector distance coincide. **(D)** The percentage of the vector distance (feeder to hive in arena 1) traveled during homeward paths in arena 2 for all three displacement conditions. Bars: medians; boxes: lower and upper quartiles; whiskers: sample minima and maxima; points: outliers; horizontal dashed lines: vector distance (all conditions) and the distance between displacement location and fictive hive entrance (edge displacements). Shaded regions as in (C).

To conclusively demonstrate that *B. terrestris* were indeed homing using a path integration vector, we observed homeward paths of foraging animals after passively displacing them to a second arena. Bees were individually allowed to exit the hive and locate the feeder, where they were trapped. The closed feeder containing the bee was moved to a second, identical arena that lacked the hive entrance. The bees were either displaced to the center of the second arena (the same position it would be in the first arena) or to a position near the edge of the second arena. For edge-displacements, a path integration-based home vector should direct the bees to the fictive hive’s position at the arena center. Therefore, the lengths of homeward paths from both center-and edge-displacement experiments should be equal, both representing the length of the home vector (Figure 3A). Conversely, if the bees navigate to the position corresponding to that of the actual hive in the first arena after edge displacements, they would have to be locating their hives by an alternate strategy, for instance by visually matching their position to a unique visual environment at the hive’s location.

In all experiments, the bees’ behavior was consistent with a path integration-based strategy. When they were displaced to the second arena, homeward paths lengths from both edge-and center-displacement experiments were close to the beeline distance from the hive to the feeder in the first arena (Center: 90.6% ± 6.4% of beeline distance, Edge: 104.2% ± 13.1% of beeline distance (Mean ± Standard Deviation); Figure 3C, D). As predicted, for the edge displacement experiments, the endpoint of the home vector (indicated by the initiation of search behavior) was located close to the center of the second aren a (Figure 3C, center panel, and Video S2).

When one of the two potential displacement locations were not in use, a fitted plug sealed the hole in the arena where the feeder could be fit. In order to control for any effect the small crack between the plug and the base of the arena may have had on influencing a homing bee to end its home-vector and start a search pattern, edge displacement experiments were repeated when the polarized light field over the second arena was offset 45° from the polarized light field in the first arena, orienting homing bees away from the center of the arena and the plug that was located there (for detailed analysis of polarization orientation, see results below). In this condition, most bees followed the rotation of the polarizer and produced home vectors similar in length to the beeline distance from the hive to the feeder in the first arena (100.7% ± 35.5% of beeline distance (Mean ± Standard Deviation); Figure 3C, D). Taken together, the results from our displacement experiments strongly support the hypothesis that foraging bumblebees use path integration while walking.

While path integration most easily explains the consistent lengths of the homeward paths, it cannot explain all directional choices the bees made. We noted that bees did not orient towards the wall of the arena during the first edge-displacement experiment, so that the expected bimodal orientation pattern became unimodally directed towards the fictive nest. Similarly, during edge-displacement experiments with a 45° polarizer offset, some bees were biased away from the arena wall closest to where the fictive nest should have been located according to a path integration home vector in that situation. These observations indicate that the proximity to the walls (and likely the visual stimuli they offer) additionally influences the homeward navigation of bees in the arena. Nonetheless, following path integration-based vector information is clearly the dominant navigation strategy bees utilized during our experiments.

### Bumblebees orient using overhead polarization patterns while walking

Path integration requires that an animal possess a mechanism to determine its orientation and an odometer to measure the distances it travels. Since *B. terrestris* were able to orient correctly towards their hive during homing, we next tested if the bees were orienting using the overhead polarization field we provided, as the bimodal distribution of homewards directions from the feeder in the above experiments and orientations during displacement experiments with a 45° polarizer offset suggested.

Bees were allowed to freely forage under a polarized light field for three days. Following familiarization, once a bee located the central feeder, it was trapped and the polarized filter was rotated 90° from its original position. In this condition, if a bee oriented using the overhead polarization field, homeward paths should be oriented perpendicular to the direction of the hive (Figure 4A). In control trials, the polarizer remained in same position between outbound and homeward paths. During these experiments, homeward paths should be oriented towards or opposite to the hive’s location (since a polarized light pattern only offers an axis of orientation). When the polarized field was static, homeward paths were oriented parallel to the direction of the burrow (data were doubled to create a unimodal distribution (see methods), 348.05° ± 40.2°, P < 0.001; Figure 4B; all orientation statistics are summarized in Tables S1 and S2). When the polarized field was rotated 90°, individuals oriented their homeward paths perpendicular to the direction of the burrow (doubled data, 161.75° ± 32.6°, P < 0.001; Figure 4B and Video S3).

**Figure 4.**
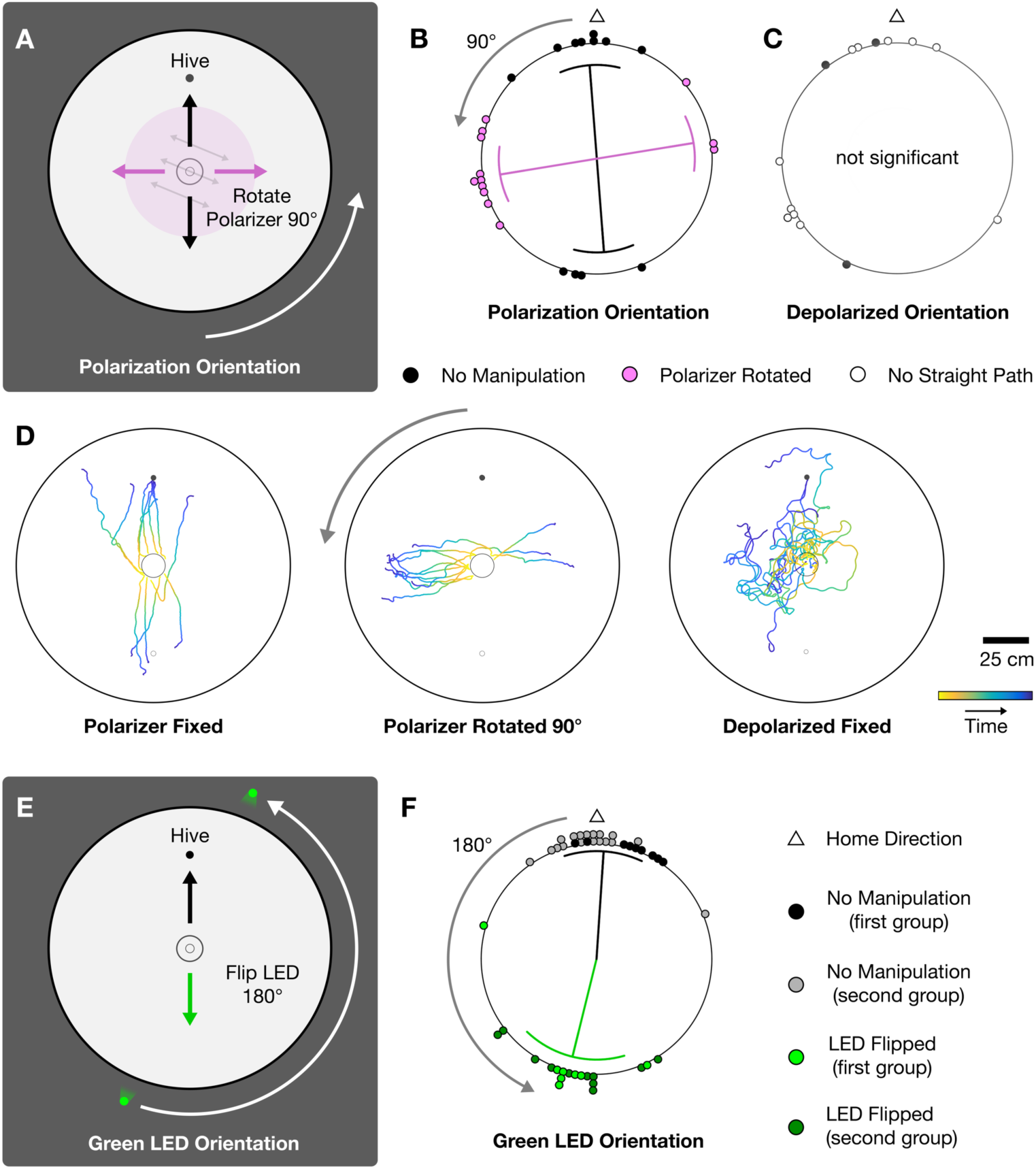
*Bombus terrestris* use artificial celestial cues present in the arena for orientation during path integration while walking. **(A)** Polarization orientation experimental design: An overhead polarized light field was rotated 90° from its original position once animals reached the feeder. Homeward paths were predicted to be oriented perpendicular to the direction of the hive if overhead polarization patterns were used for orientation. **(B)** Orientations of homeward paths for polarization orientation experiments. The polarized light field was either fixed in place or rotated (direction: curved grey arrow). Arrows in each plot: mean vectors (arrow angles: circular mean, curved arrowheads: circular standard deviations; arrow lengths: strength of orientation 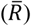). Both groups exhibited significant orientations and were significantly differently oriented from one another (p<0.05). **(C)** Orientations of homeward paths under a depolarized light field. Homeward orientations were not significantly oriented (p=0.53). **(D)** Path tracings from all homeward paths before search behaviors were initiated under polarized and depolarized light. Straight homeward paths were not observed under depolarized light fields. **(E)** Point source orientation experimental design: A green LED point source was flipped 180° from its original position once animals reached the feeder. Homeward paths were predicted to be oriented in the opposite direction of the hive if *B. terrestris* were orienting using the point source. **(F)** Orientations of homeward paths for point source orientation experiments. All groups exhibited significant orientations and manipulated groups were significantly differently oriented from not manipulated groups (p<0.05). For summary of all orientation statistics see Tables S1 and S2.

To ensure bees were indeed orienting to the polarization pattern the filter created rather than other unintended artifacts the filter may have created, such as potential small differences in brightness across the filter, control experiments were conducted during which the filter was configured to provide a depolarized light field. Under this condition, bees no longer exhibited straight homeward paths, and these paths exhibited no significant orientation (P = 0.526, Figure 4C, D). Taken together, these results show that *B. terrestris* use overhead polarization information when available to orient while walking.

### Bumblebees orient using a visual point source while walking

Besides the skylight polarization pattern, the Sun itself is a prominent directional cue that many animals use for orientation in nature. Thus, we next determined if bumblebees would use a bright point source for orientation if provided instead of an overhead polarization field. A high-powered green-LED was set at an azimuth of 157.5° from the hive entrance with an elevation of 21.8° when viewed from the center of the arena. Following three days of familiarization in this new setting, bees were individually allowed entrance to the arena. When they had reached the central feeder, they were trapped, the LED was turned off and was replaced with an identical LED 180° from the original’s position. If the bees orient using the point source provided, homeward paths should be oriented away from the hive in this condition (Figure 4E). In control trials, the LED remained at the same location during outbound and homeward paths.

During experiments when the position of the LED was flipped 180°, homeward paths were primarily oriented in the opposite direction of the hive (197.45° ± 34.15°, P = 0.001). During control trials when the LED was left in place, homeward paths were oriented towards the hive (17.36° ± 14.55°, P < 0.001; Figure 4F and Video S4). Since the homeward directions from the first set of these point sour ce experiments appeared clustered directly opposite of the green LED rather than around the expected position of the hive, we examined whether this bias was consistent in a repetition of the experiment with independently trained bees. Orientations from the second set of trials were more clearly centered towards or away from the direction of the hive in the predicted manner rather than away from the direction of the LED (fixed: 356.83° ± 20.43°, P < 0.001; rotated: 191.38° ± 26.07°, P < 0.001; Figure 4F), suggesting that the bees perform true menotactic orientation rather than simple negative phototaxis. Overall, these results indicate that *B. terrestris* can orient using a bright point source while walking, possibly interpreting the bright LED as a Sun cue.

### Bumblebees prefer to orient using a point source to overhead polarization patterns

Similar to our bumblebees, many animals can use more than a single celestial cue for orientation. When orientation responses are examined in a natural environment or in a laboratory setting, some cues are weighted more heavily than others and therefore dominate in experimentally generated cue conflict situations^6,12,14,20,38^. Since we demonstrated that *B. terrestris* will orient to overhead polarization patterns and a point source when both cues were in isolation, we next were interested in which of the two cues the bumblebees preferred to orient with when both were available in concert.

As in previous experiments, bumblebees were allowed to freely forage in the arena for three days while the overhead polarization field and the point source were both present in the arena. These cues were arranged so that the bright LED was located at 90° with respect to the angle of polarized light at an elevation of 21.8°, loosely mimicking the natural skylight situation close to dusk or dawn. After familiarization, bees were allowed to forage individually. Once a bee was trapped in the feeder, both orientation cues were rotated by 90°. Considering the results from the previous experiments, bees were predicted to orient 90° from the direction of the hive in this condition. Finally, each orientation cue was individually rotated by 90° while the other cue remained in its original position. If *B. terrestris* weights one of the two cues more strongly, the bees should follow the stimulus rotation only when the dominant cue had been rotated (Figure 5A).

**Figure 5.**
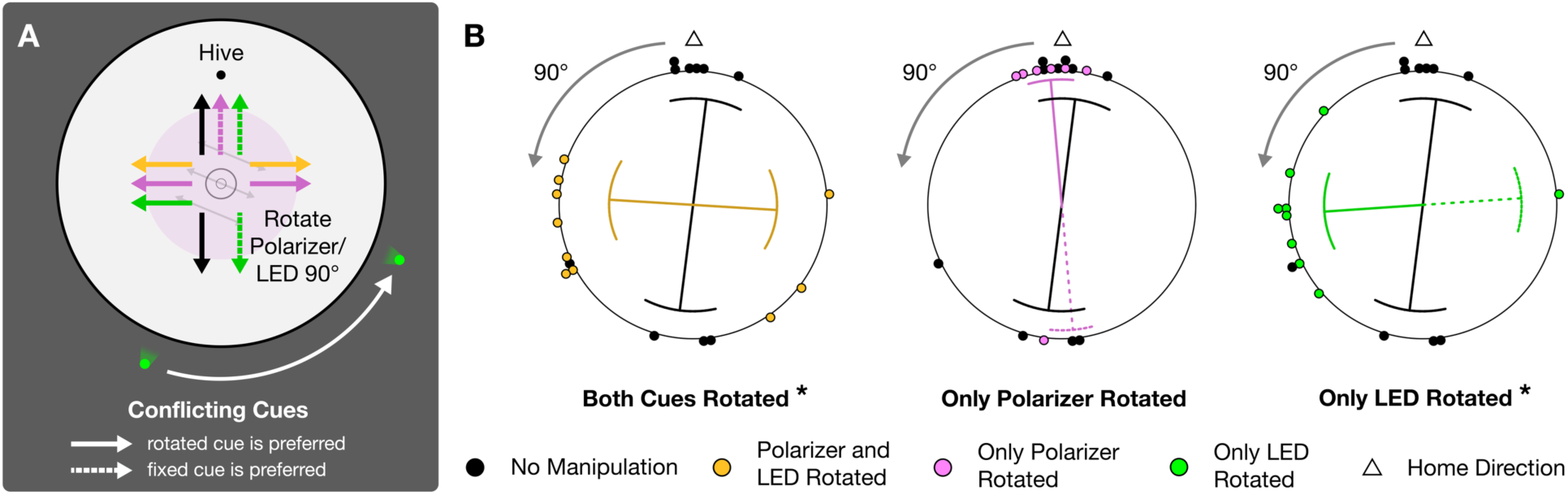
*Bombus terrestris* prefer to orient using a point source rather than an overhead polarization field in the arena while path-integrating. **(A)** Conflicting cues experimental design: To determine if bumblebees preferred to use the polarized light field or the green LED for orientation when both are present in the arena, both cues were rotated 90° either together or in isolation. If the bees prefer one of the two orientation cues, they should only orient 90° from the hive when the dominant cue has been rotated. **(B)** Orientations of homeward paths for conflicting cue orientation experiments. Under each condition, the orientation cues were either fixed in place or rotated (direction: curved grey arrows). All groups exhibited significant orientations (p<0.05). Asterisks (*) indicate manipulation conditions with group orientations that were significantly differently oriented from trials when no manipulations occurred (p<0.01). Arrows in each plot: mean vectors (arrow angles: circular mean, curved arrowheads: circular standard deviations; arrow lengths: strength of orientation 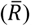). For summary of all orientation statistics see Tables S1 and S2.

When neither cue was manipulated, bees oriented significantly in the axis of the hive (doubled data, 7.39° ± 38.96°, P < 0.001). During experiments in which both cues were rotated, bees oriented perpendicular to the hive direction, as expected (doubled data, 181.52° ± 55.41°, P = 0.016). During trials when the orientation cues were rotated in isolation, bees ignored the rotation and oriented towards the hive when only the polarization pattern was rotated (doubled data, 349.8° ± 20.63°, P < 0.001), and followed the stimulus manipulation when only the point source was moved (doubled data, 169.31° ± 44.69°, P = 0.004). These results demonstrate that bees weighted the point source more highly than the polarization field in the navigation arena and consequently ignored the polarization stimulus when choosing a walking direction if it conflicted with the azimuth information of the point source (Figure 5B).

While our data provides a clear cue hierarchy for the specific stimulus conditions of our arena, the artificial nature of the provided orientation cues limit generalizations to a natural environment. Both absolute and relative light intensities for natural celestial cues differ from the artificial cues in the arena. Although orientation behaviors should be robust to such changes given that these intensities vary highly in nature as well, cue preferences in bumblebees might change under more naturalistic conditions. Nonetheless, matching our results in bumblebees, honeybees interpret a long-wavelength light spot as the Sun^39^ and can also use a strong artificial polarization field to orient their waggle dances, communicating their previous foraging paths away from the hive under the natural sky^12^. Also, von Frisch^12^ found that when the Sun and a small overhead polarized light cue were placed in conflict, honeybees will orient their dances solely to the Sun. However, the influence of the polarizer became stronger with increased amounts of UV light transmitted. Celestial cue preferences in honeybees thus depend on the relative strengths of the two cues^40^, a finding that is likely to also be true for orientation cue preferences in bumblebees.

### Bumblebees perform systematic search patterns when home-vectors fail

In the previous experiments, the end of a home run of a bee was defined by a sudden change in extended walking direction, revealing the bee’s estimate of where its hive should be. However, this position often did not coincide with the actual entrance of the hive. If that was the case, the bees executed stereotyped search behaviors to locate their home, a key feature of path integration in many species. In most experiments, these searches were located around a nest position that was close to one of the arena walls, so that the resulting search pattern was geometrically biased by the limits of the arena. We therefore analyzed search behaviors only from the edge-displacement experiments under a polarized light field that was not offset between the two arenas. In those conditions, searches started near the center of the arena and therefore were geometrically unbiased. These observed search behaviors were composed of loops that started and ended near the search initiation point, therefore being most focused around the search origin, and increased in size over time (Figure 6A, B).

**Figure 6.**
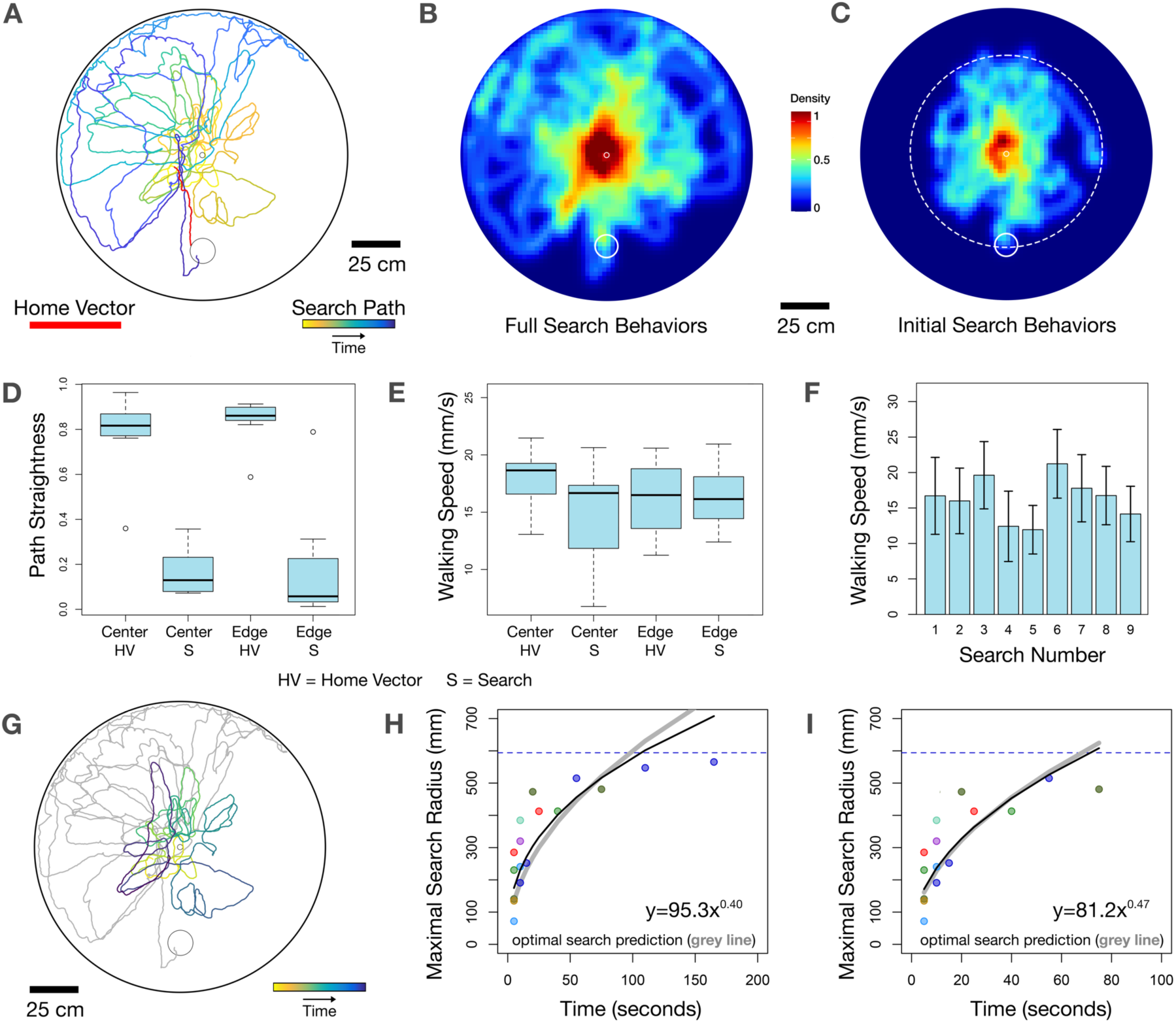
*Bombus terrestris* perform systematic search behaviors when path integration home vectors fail to locate the hive. **(A)** Example of a search behavior during a trial when a bee was displaced to the edge of arena 2. Larger empty circle: displaced feeder; Smaller empty circle: vector position of the fictive burrow. **(B)** A heat map of all search behaviors compiled during edge displacement experiments with no polarizer offset. **(C)** A heat map of all search behaviors in (B) up to a distance of 25 cm from to the arena wall (dashed circle), used for analysis in (H). **(D)** Straightness of paths during the homing and search phases of center-and edge-displacement experiments (where path straightness = beeline distance/path length). Home-vector paths were straighter than search paths (p<0.001; all other pairwise comparisons, p>0.1). **(E)** Walking speed remained approximately constant during homing and search phases (p=0.512; χ^2^=3.284). Bars: medians; boxes: lower and upper quartiles; whiskers: sample minima and maxima; points: outliers. **(F)** Mean walking speed of each bumblebee during the search phase of edge-displacement experiments. Bars show average walking speeds over one second intervals throughout a search for each bee, with error bars (± SD) indicating relatively constant walking speeds during searches. **(G)** Search behavior from (A), but with the colored section highlighting completed loops before the search expanded to a distance of 25 cm to the wall, as used for analyses in (H). **(H)** Absolute radii of searches in (C) with at least one completed loop before the search expanded to a distance of 25 cm from the arena wall (n = 8). The maximum distance of each increasing search loop from the search origin is plotted against time after search initiation. Colors: individual searches. Black line: the power function of best fit for all data, yielding a search expansion relation with radius_max_ ∝ time^0.4^. Optimal search theory predicts that searches should expand by radius_max_ ∝ time^0.5^ (grey line). Dashed blue line: the distance between the search origin and the 25 cm limit from the arena wall for the search plotted in the same color. **(I)** Same as (H) except constrained to the first 100 seconds of search. The initial search expansion (radius_max_ ∝ time^0.47^) better fits the expansion predicted by optimal search theory. Also see Figure S2.

In other path integrating animals, homing usually consists of a straight, quick home vector followed by a contorted and often slower search pattern^3,29,41^. In contrast, we found that even through bumblebees exhibited search patterns that were much more tortuous than home vectors, walking speed remained relatively constant throughout both phases of homeward travel and over the duration of the search (Figures 6D-F).

To examine whether walking bumblebee search patterns quantitatively resemble those observed in other species as well as those predicted by optimal search theory, we determined the rate of the bees’ search expansion by measuring the farthest distance between the bee and the search initiation point during each successive increasing loop in the search, until bees reached a distance of 25 cm to the arena wall (the distance of the hive entrance from the arena wall). This limit was imposed to prevent the constraints of the arena wall to impact the free expansion of the search. Assuming constant detection width of the nest, constant speed, and a Gaussian uncertainty distribution, optimal search theory predicts a parabolic shape of the search distribution to maximize search success. This results in an expected increase of search radius proportional to the square root of the search time, radiusmax ∝ time^0.5 1^. Data from desert ant searches^27^ match this prediction, with the maximum search radius being proportional to time^0.48 1^. Similarly, the radii of searches in mantis shrimp following path integration home vectors expand at a rate proportional to time^0.43 29^. By fitting a power function to our data, we observed that the search patterns of walking bumblebees expand at a rate proportional to time^0.40^ (Figures 6C, G, H, and S2). When only the initial expansion of the search is observed, further limiting the potential constraining effect of the arena wall, the expansion rate of the searches was proportional to time^0.47^ (Figure 6I), very closely y approximating the expansion of searches predicted by optimal search theory and matching those exhibited by other path-integrating animals.

## DISCUSSION

Our results are the first to conclusively demonstrate that bumblebees use path integration to navigate and is one of the first to show that bees path integrate over short distances while walking. Previous work on navigation in walking honeybees^42^ found that bees forced to walk performed waggle dances very similar to those of flying bees. These walking bees transitioned from round dances (indicating feeders at short distances) to waggle dances (indicating feeders at further distances) at a much earlier stage (3-4 m rather than 50-100m). Thus, walking foraging resembled a scaled down version of foraging during flight in key aspects. Importantly, when honeybees were forced to approach and return from a feeder along a curved or 90° angled path, the waggle dances they performed back in the hive correctly indicated the bee-line direction of the feeder, suggesting that the walking bees performed path integration during the outbound trip. Reflecting these results in walking honeybees, we found that in the laboratory, walking bumblebees path-integrate by using polarized light fields and point sources as directi onal cues. They also estimate vector distances accurately, yet it remains unclear what mechanism they use to track distances in our arena. Stride integration, optic flow, or a combination of the two are likely contenders. Walking bumblebees indicate the arrival at their estimated hive position by performing stereotyped search patterns that both qualitatively and quantitatively resemble those found in other path integrating species, like ants. Taken together, the ability to path integrate while walking appears to be a feature present across a wide range of hymenopterans, including species that usually fly.

While our assay provides a novel and robust paradigm to dissect path integration, the question arises of how biologically relevant those refined path integration abilities over short distances in walking bumblebees are, as these animals typically fly over long distances while foraging. Despite their aerial lifestyle, bumblebees almost exclusively walk in the hive and will also occasionally walk over short distances while away from the hive in nature^43^. Some species, as reported for an Amazonian bumblebee (*Bombus transversalis*), even build extensive walking trails around their nest and thus resort to a relatively terrestrial lifestyle^44^. Chittka et al.^30^ found that bumblebees (*Bombus impatiens*) also choose to walk rather than fly when forced to forage in darkness. During this study, bumblebees could walk to a feeder but were unable to make oriented homeward paths after feeding, similar to our experiments under a depolarized light field. As suggested by modelling studies^1,45–47^, path integration relying on purely idiothetic (self-motion) directional cues likely accumulates too much error to allow for successful homeward navigation in situations where clear environmental directional information is absent. In each case, bees must maintain a stable navigational sense while walking, in addition to their well-known ability to navigate while flying over long distances. Path integration offers a solution to that need and could be implemented by swapping the potential sources of distance information, e.g. between optic flow (used during flight) and a stride integrator (used during walking), thereby co-opting the same neural machinery for flight and walking.

The behavioral paradigm established here can be used to test many navigationally related behaviors beyond what we have described, including mechanisms of odometry during walking. It also offers the chance to dissect more sophisticated strategies, including the generation of food vectors and novel shortcuts, landmark learning, integration of landmark and vector navigation, and developing optimal routes while foraging from multiple places (for hypotheses of how insects may neurally accomplish some of these tasks by using vector navigation, see Le Moël et al.^48^). Further, our behavioral system can serve as a powerful comparative tool between animals that utilize similar navigation strategies. For example, mantis shrimp foraging underwater have been shown to exhibit strikingly similar path integration behaviors in comparably-sized arenas to what we present here in walking bumblebees^6,29^. Comparative behavioral studies across a wide range of species promise insights into how similar navigational problems are solved by evolutionary disparate animals navigating in different media.

Finally, our paradigm offers a clear path towards probing the neural basis of path integration during ongoing behavior. Over recent years, understanding of the neural basis of navigation behaviors in insects has increased substantially due to detailed functional studies and neuroanatomical descriptions of navigationally relevant brain regions. From these studies, the central complex, a conserved set of arthropod brain regions, has emerged as the likely navigational control center that both integrates navigationally relevant stimuli^35,49–55^ and directs steering movements during navigation^56^. By combining our behavioral arena with electrophysiological techniques used to record from either freely behaving animals^56^ or by transferring our paradigm to a virtual reality setup for tethered behavior^41,51,53–55,57^, hypotheses regarding the underlying neural mechanisms for path integration vector memory formation^35^ can be directly tested.

## Acknowledgements

This research has received funding from the European Union’s Horizon 2020 research and innovation programme under the Marie Skłodowska-Curie grant agreement No. 101027405 (to R.N.P.) and the European Research Council under grant agreement no. 714599 (to S.H.), as well as the Swedish Research Council (2018–04851, to S.H.). We thank Allen Cheung for his insight into optimal search theory and advice for our search analyses. We are also grateful to Johan Bäckman (LU Biology electronics workshop) for technical assistance in building the arena lighting system and Lars Fredricksson (LU Biology mechanical workshop) for help with planning and constructing the behavioral arena and Nils Sundqvist for additional assistance. Finally, we thank all members of the Heinze lab and the Lund Vision Group for helpful discussions.

## Author Contributions

Conceptualization, R.N.P. and S.H.; Methodology, R.N.P.; Investigation, R.N.P. and J.K.; Analysis, R.N.P. and J.K.; Writing-Original Draft, R.N.P.; Writing-Review & Editing, R.N.P. and S.H.; Funding Acquisition, R.N.P. and S.H.; Guidance, S.H.

## METHODS

### Lead contact and materials availability

Further information or materials associated with this research will be made available upon reasonable request to the lead contact author, Rickesh Nitin Patel. This study did not generate any new or unique reagents.

### Experimental model and subject details

Commercial hives of *Bombus terrestris* (Natupol Smart, Koppert Biological Systems) were shipped to Lund University and were housed at room temperature under a 12:12 light:dark cycle. Two days after arrival, the wings of all individuals were clipped. One day following clipping, the hive was introduced into the experimental arena and bees were allowed to freely forage. They were allowed access to a feeder with 1M sugar-water solution and ground pollen (Bipollen EKO, Rawpowder Sweden AB) placed at the center of the arena. Foraging bees were labelled with unique small colored tags adhered to the back of the thorax. Data were collected from 62 individual bees from three hives, with each individual only being used once per experiment.

### Method details

#### Experimental Arenas

Two relatively featureless, 1.5 m-diameter circular navigation arenas were constructed with a white composite wood base and were elevated 1.2 m to allow access below the arena (Figures 1A, B and S1). The first arena contained a hive entrance 25 cm from the arena’s periphery. The hive was placed beneath the arena and was connected to the arena by a tube with three doors to have control of bees that wished to enter the arena (Figure S1D). Sugar-water and pollen were placed in a removable conical feeder at the center of the arena. The conical feeder had a door which could slide over the bee while it was feeding, trapping it in the feeder (Figure S1C). Both the feeder and the hive-entrance were placed below ground-level of the arena so they were not visually detectable by bumblebees while in the arena. Trials were recorded from above using a raspberry pi camera connected to a raspberry pi unit (Raspberry Pi Foundation) outside of the arena, where experiments were observed and trials recorded.

The second arena was identical to the first with the exception that no hive or hive entrance was present and two potential locations were present where a conical feeder could be placed (at the center and near the edge of the arena). When a feeder was not occupying one of these two potential feeder locations, a tight plug, level with the arena floor and constructed of the same material as the base of the arena was present instead.

Arenas were placed in a dark room and surrounded by thick matte black curtains and lit from above using a centered, diffused, custom built LED light source with sets of ultraviolet (40 units, LZ1-00UV0R, LED Engin Inc.) and white LEDs (37 units, LZ1-10CW02, LED Engin Inc.). Composite filters constructed of a linear polarizer (38% transmission neutral grey, Rosco Laboratories Inc.) and two sheets of tracing paper were placed under each light source. When the polarizer side of the composite sheet faced downwards towards the arena, light was linearly polarized to an average degree of 99.78% from 350 to 600 nm (98.23% from 350 to 450 nm). For unpolarized fields, the depolarizing waxed paper side faced downwards, reducing the average degree of polarization to 2.32% from 350 to 650 nm (1.85% from 350 to 450 nm) (Figure 1D). The overhead polarization stimulus had an angular diameter of 22.6° of when viewed at ground level from the center of the arena and could be rotated up to 90° from outside of the arena. Four green LED (LZ4-00G108, LED Engin Inc.) point sources were positioned at 90° intervals around the arena with one at a 22.5° offset from the direction of the hive entrance from the center of the arena at an elevation of 21.8° above ground level when viewed from the center of the arena. All four LEDs could be controlled from outside the arena. Since the long-wavelength photoreceptor spectral-class in bumblebees provides achromatic contrast^37^, green LEDs were chosen to create a bright point source. Irradiance spectra of all light sources can be reviewed in Figure 1C.

#### Spectrometry

Irradiance measurements were taken from the center of the arenas at ground level using an Ocean Optics USB2000 spectrometer (Ocean Optics Inc.) connected to a 400 µm diameter, fiber-optic cable with a cosine-correcting head. Irradiance measurements of the green LED point source were taken with the cosine-corrector angled towards the light source at a distance 20 cm from it (Figure 1C).

Transmittance measurements of the polarizing filter were taken using the same spectrometer system without a cosine-correcting head. The percent polarization of light transmitted through the filter was determined by calculating the ratio of the percent transmission of the polarizing filter layered with a second polarizer oriented first perpendicularly and then parallel to the first filter, both with the polarizing and depolarizing side of the filter facing upwards (Figure 1D).

#### Hive Observations

A hive containing bees with clipped wings were introduced into the experimental arena and were allowed to freely forage with a polarized light source present overhead and a feeder with 1M sugar-water and pollen at the center of the arena. During the first day, each individual bee that emerged from the hive was labeled with colored tag. During the following two days, the number of times each bee made a foraging trip from the hive to the feeder and back within the hour was recorded over the course of the full 12-hour light-cycle. The foraging activity of the hive was recorded one day per week for an additional three weeks following the initial two observation days. Any bees emerging from the hive without an identity tag following the initial week had their wings clipped, were given a tag, and were assigned to a new cohort.

#### Experimental Procedures

A hive containing bees with clipped wings were introduced into the experimental arena and were allowed to freely forage for three days. During familiarization, a feeder with sugar-water and pollen was placed at the center of the arena. During this period, either the overhead polarized light field (before polarization orientation and displacement experiments), the green LED point source (before green point-source orientation experiments), or both light sources (before conflict experiments) were present.

After familiarization, the doors to the hive were closed and experiments were conducted. During experiments, bees were let out of the hive individually, allowed to locate the feeder, and were trapped in the feeder. During this time, any experimental manipulations were performed before the bees were released and allowed to return to the hive. All conditions per experiment were performed in a randomized order.

During experiments in which animals were displaced, once bees were trapped in the feeder, the entire feeder was removed from the underside of arena 1 and placed at either the center or edge position in arena 2. Bees were then released and allowed to leave the feeder. Once they returned back to the feeder, they were trapped in the feeder once more, displaced back to arena 1, and allowed to return back to the hive (Figure 3A). To control for any effect the edge of the plug filling the center feeder position during edge-displacement experiments may have had on influencing a homing bee to end its home-vector and start a search pattern, edge displacement experiments were repeated when the polarized light field over the second arena was offset 45° from the polarized light field in the first arena, orienting homing bees away from the center of arena and the plug that was located there.

During experiments in which the polarization field was manipulated, once an experimental animal found the central feeder, the bee was trapped and the composite polarization filter with the polarizer-side facing downwards over the arena, creating a linearly polarized field, was rotated 90° from its original position and the bee was released. In control trials, the polarizer was rotated 45° forward and backwards by the researcher, so it ended at its original position (Figure 4A). To verify that bees were orienting to the polarization pattern the filter produced, the filter was placed diffuser-side downward over the arena, creating a depolarized field during the entire trial and was rotated 45° forward and backwards so it ended at its original position.

During green point-source orientation experiments, trials were run in the presence of a single green LED 157.5° relative to the hive entrance from the center of the arena. Once a bee was trapped at the central feeder, the green LED was turned off and an identical one was turned on 180° from the original LED’s position and the bee was released. As a control, the green LED was turned off and turned back on at its original position when a bee was trapped at the feeder.

During conflict experiments, both the overhead polarization field and the green LED was present. Once an animal was trapped at the feeder, one of four conditions were enacted: 1. The polarizer was rotated 45° forward and backwards and the green LED was turned off and back on at the original position (neither cue’s position was changed). 2. Both cues were rotated 90° in the same direction. 3. The polarizer was rotated 90° and the green LED remained in the same position. 4. The green LED was rotated 90° and the polarizer remained in the same position.

### Quantification and statistical analysis

Foraging paths to and from food locations to the hive were video recorded at 10 frames per second. Videos were unwarped to correct for lens distortion and distortion due to camera placement using the Camera Calibration Toolbox in Matlab (vR2020a, MathWorks Inc.). To differentiate homeward paths from continued arena exploration, paths from the feeder were considered to be homeward paths when a straight path started at most 14 cm from the feeder and when they did not deviate more than 90° from their initial trajectories for at least 28 cm. From these homeward paths, search behaviors were determined to be initiated when an animal turned more than 90° from its initial trajectory and continued walking at a distance of at least 14 cm.

Paths were automatically tracked using the software ivTrace (v2.23, Bielefeld University), from which the output is given as Cartesian coordinates. From these measures, home vector lengths and beeline distances from the feeder to the hive entrance were determined. The straightness of paths and mean walking speeds during both the home vector and search phases of homeward travel were also calculated. Further, for search behaviors from edge displacement experiments without an offset in polarizer orientation between the two arenas, mean walking speeds binned over one second increments (10 frames) were calculated for the entirety of the search.

Additionally, the orientation of homeward paths when bees were 28 cm from the feeder in relation to the hive entrance was recorded using the angle tool in ImageJ (v2.3.0, Fiji).

Search behaviors with at least one completed loop were analyzed from all trials when animals were displaced to the edge position of arena 2 with the polarizer set in the same position between the two arenas (n=8), since searches should start at approximately the center of the arena, negating the potential effect the proximity to the arena wall would have on biasing the geometry of the search. The radii of these search behaviors were measured as the farthest distance of each successive increasing search loop from the point of search initiation (i.e. the end point of the home vector) using the line tool in ImageJ. To limit the potential effect the proximity to the arena wall may have had on search expansion, the absolute radius of each search loop was measured from the beginning of the search until bees reached a distance of 25 cm from the arena wall. To observe the general expansion pattern of the maximal search radius, only search loops larger than previous loops were included. These values were plotted over time and fitted with a power function. To remove the potentially distorting impact of a single, very long search, this analysis was repeated for only the initial 100 seconds of each search. This further limited the influence of the arena wall on free search expansion.

All statistical analyses were run on R (v4.0.2, R Core Development Team) with the “autoimage”, “CircStats”, “circular”, “ggplot2”, “plotrix”, and “shape” plugins. All statistical analyses for experiments conducted under a polarized light field were performed after using the doubling angles procedure for bimodal data outlined in Batschelet^58^. Orientation data were analyzed using the following procedures for circular statistics^58^: Rayleigh tests of uniformity were used to determine if homeward paths were oriented within a group for all experiments. Watson Two-Sample tests for homogeneity were used to determine if group orientations were significantly different from one another. All reported mean values for orientation data are circular means and circular standard deviations.

Kruskal-Wallis tests were used to evaluate differences in the path straightness and walking velocity during the home-vector and search phases of homeward paths during displacement experiments. If significant differences were observed, post-hoc pairwise Wilcoxon ranked tests were used to determine which specific groups were different from one another.

Bonferroni corrections were used for all tests when applicable. All statistical information including sample sizes, test statistics, P-values, means, and standard deviations are presented in Tables S1 and S2.

## SUPPLEMENTAL INFORMATION

**Supplemental Figure 1.**
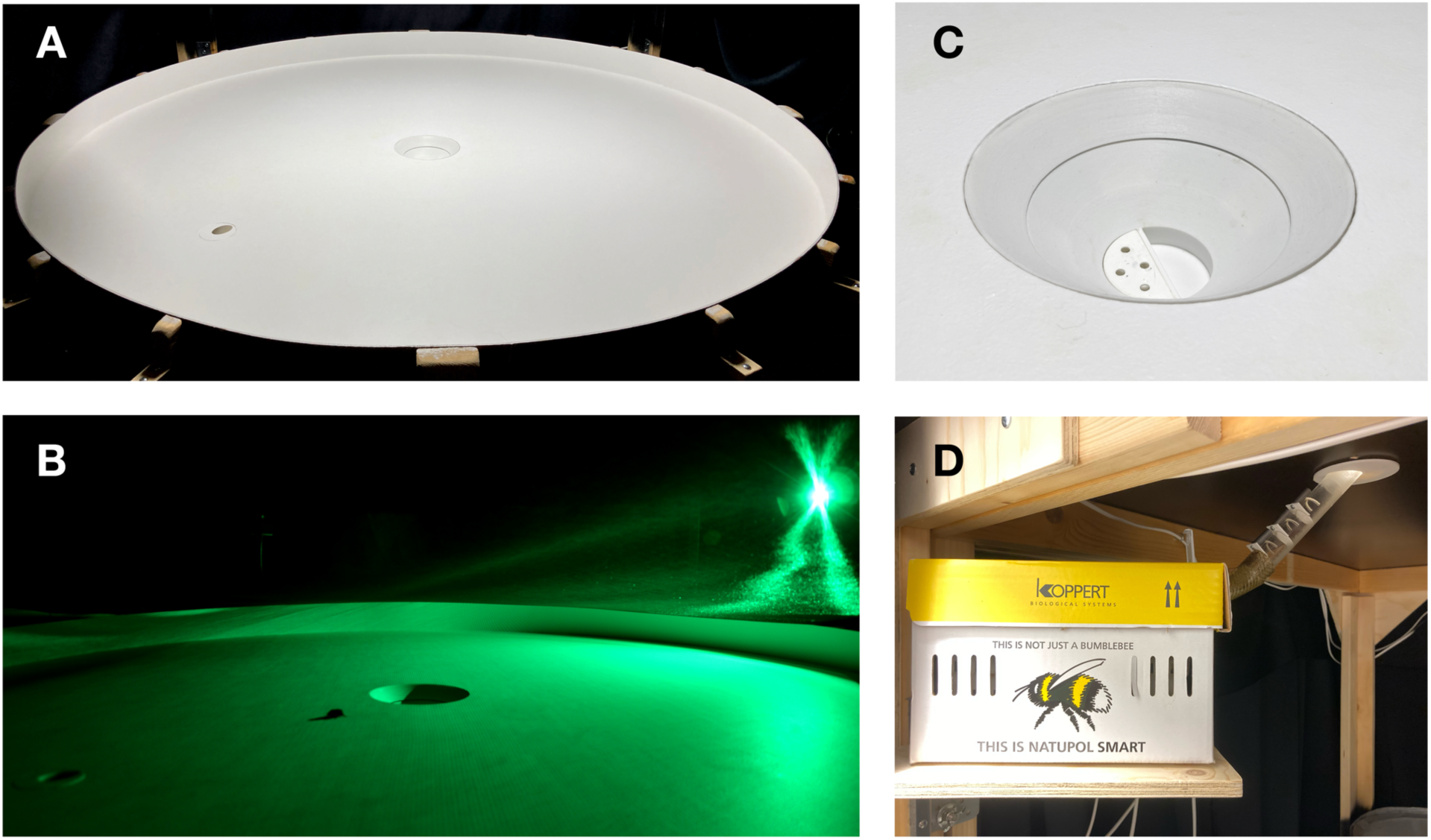
Arena details. Photos of the behavioral arena illuminated by **(A)** the overhead UV and white LEDs transmitted through the polarized filter and **(B)** the green LED point source. **(C)** Photo of the feeder at the center of the arena viewed at an angle from above. Pollen and sugar water may be placed in the center of the feeder. A door can be slid through the thin slit on edge of the feeder, trapping a feeding bee. The entire feeder can be removed from the underside of the arena. **(D)** Photo of a commercial hive of bumblebees connected to the underside of the arena by a tube with doors. All three doors are in a closed position. Related to Figure 1.

**Supplemental Figure 2.**
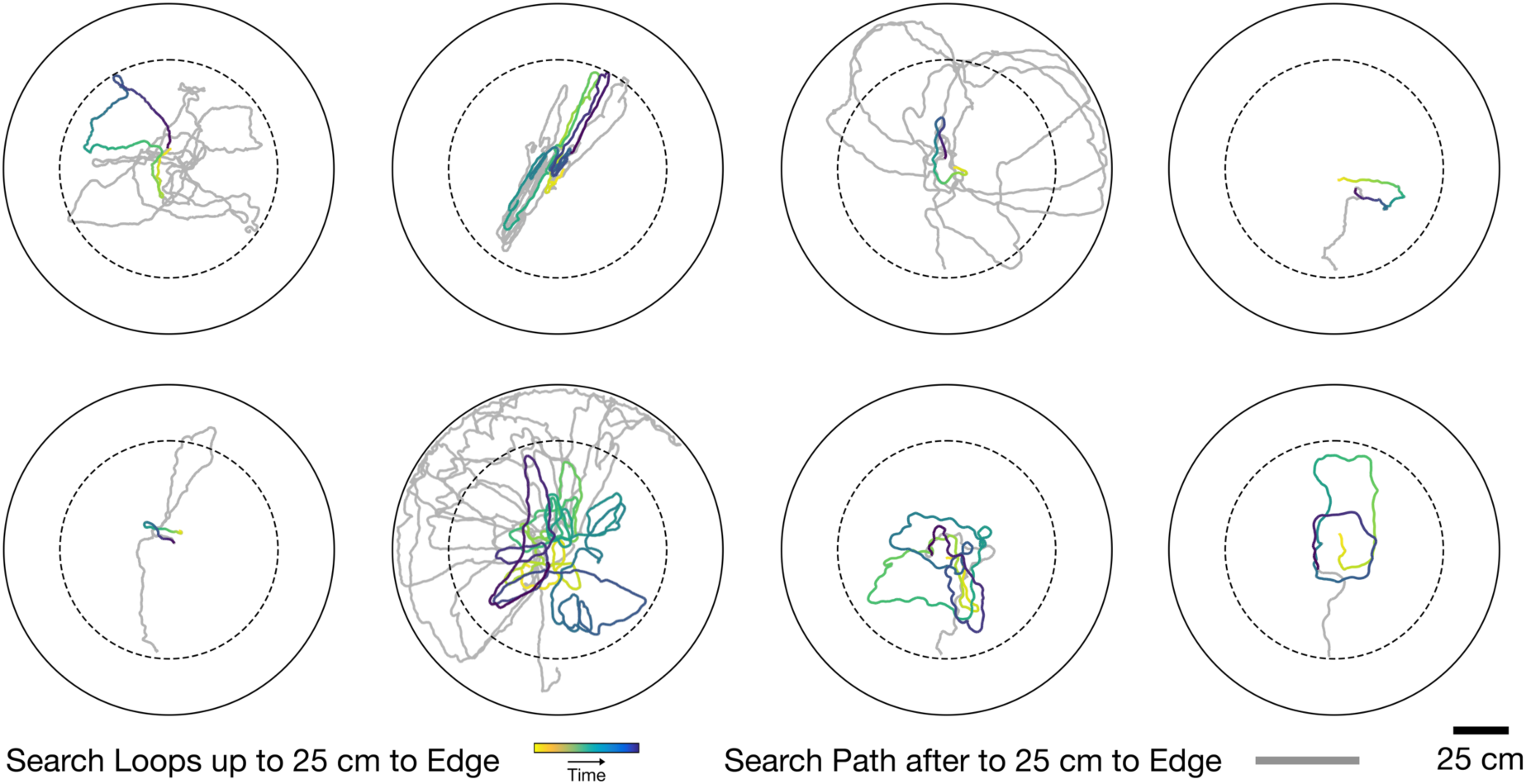
Search behaviors from edge displacement experiments without a polarizer offset between the two arenas. Eight search behaviors contained at least one completed loop until the searches expanded to a distance of 25 cm from the arena wall (dashed circle). These completed loops are presented in a colored gradient. The remainder of the searches are in grey. Related to Figure 6.

**Supplemental Table 1.** Summary of orientation statistics for all experimental groups. Orientations were analyzed using Rayleigh Tests of Uniformity. Significance indicates that groups are oriented in a single direction. Only orientations of straight-line paths (see Methods) were used for analyses. *Analyzed using doubled data. Related to Figures 3 and 4

| Experiment | P-value | N | $\bar{R}$ | Mean ± SD |
| --- | --- | --- | --- | --- |
| **Orientation to Polarized Light** |  |  |  |  |
| Polarized Field Static* | <0.001 | 12 | 0.813 | 353.26° ± 36.9° |
| Polarized Field Rotated 90°* | <0.001 | 13 | 0.850 | 161.75° ± 32.6° |
| Depolarized Field Static | 0.526 | 3 | 0.984 | NA |
| **Orientation to Point Source** |  |  |  |  |
| Green LED1 Static | <0.001 | 8 | 0.968 | 17.36° ± 14.55° |
| Green LED1 Rotated 180° | 0.001 | 9 | 0.837 | 197.45° ± 34.15° |
| Green LED2 Static | <0.001 | 18 | 0.902 | 356.83° ± 20.43° |
| Green LED2 Rotated 180° | <0.001 | 11 | 0.938 | 191.38° ± 26.07° |
| **Conflict Experiments** |  |  |  |  |
| Both Cues Static* | <0.001 | 10 | 0.795 | 7.39° ± 38.96° |
| Both Cues Rotated 90°* | 0.016 | 10 | 0.627 | 181.52° ± 55.41° |
| Polarized Field Rotated 90°* | <0.001 | 7 | 0.937 | 349.8° ± 20.63° |
| Green LED Rotated 90°* | 0.004 | 9 | 0.737 | 169.31° ± 44.69° |

**Supplemental Table 2.** Summary of Watson Two-Sample Tests of Homogeneitys. Significance indicates that groups are oriented differently from one another. *Analyzed using doubled data. ^±^Due to Bonferroni corrections, significance threshold is p<0.0167. Related to Figures 3 and 4.

| Experiment | P-value | $\sigma$ |
| --- | --- | --- |
| Polarized Field: Static vs. Rotated 90°* | <0.001 | 0.527 |
| Green LED1: Static vs. Rotated 180° | <0.01 | 0.363 |
| Green LED2: Static vs. Rotated 180° | <0.001 | 0.575 |
| Conflict: Static vs Both Rotated 90°*<sup>±</sup> | <0.01 | 0.353 |
| Conflict: Static vs Polarizer Rotated 90°*<sup>±</sup> | >0.1 | 0.061 |
| Conflict: Static vs LED Rotated 90°*<sup>±</sup> | <0.01 | 0.361 |

